# GLUT1 inhibition blocks growth of RB1-positive Triple Negative Breast Cancer

**DOI:** 10.1101/764944

**Authors:** Qin Wu, Wail ba-alawi, Genevieve Deblois, Jennifer Cruickshank, Shili Duan, Evelyne Lima-Fernandes, Jillian Haight, Seyed Ali Madani Tonekaboni, Anne-Marie Fortier, Hellen Kuasne, Trevor D. McKee, Hassan Mahmoud, Sarina Cameron, Nergiz Dogan-Artun, WenJun Chen, Ravi N. Vellanki, Stanley Zhou, Susan J. Done, Morag Park, David W. Cescon, Benjamin Haibe-Kains, Mathieu Lupien, Cheryl H. Arrowsmith

## Abstract

Triple negative breast cancer (TNBC) is a deadly form of breast cancer due to the development of resistance to chemotherapy affecting over 30% of patients. New therapeutics and companion biomarkers are urgently needed. Recognizing the elevated expression of glucose transporter 1 (GLUT1, encoded by *SLC2A1*) and associated metabolic dependencies in TNBC, we investigated the vulnerability of TNBC cell lines and patient-derived samples to GLUT1 inhibition. We report that genetic or pharmacological inhibition of GLUT1 with BAY-876 impairs the growth of a subset of TNBC cells displaying high glycolytic and lower oxidative phosphorylation (OXPHOS) rates. Pathway enrichment analysis of gene expression data implicates E2F Targets pathway activity as a surrogate of OXPHOS activity. Furthermore, the protein levels of retinoblastoma tumor suppressor (RB1) are strongly correlated with the degree of sensitivity to GLUT1 inhibition in TNBC, where RB1-negative cells are insensitive to GLUT1 inhibition. Collectively, our results highlight a strong and targetable RB1-GLUT1 metabolic axis in TNBC and warrant clinical evaluation of GLUT1 inhibition in TNBC patients stratified according to RB1 protein expression levels.

## Introduction

Breast cancer is the most common female cancer worldwide, with 1.7 million new cases and over 520,000 deaths recorded in 2012^1^. Triple negative breast cancer (TNBC) is a highly aggressive subtype of breast cancers, that lacks the expression of the oestrogen receptor α (ERα), progesterone receptors (PR) and the epidermal growth factor receptor 2 (HER2). TNBC represent 15-20% of breast cancer cases but accounts for 25% deaths^2^. In addition, TNBC has a higher metastatic rate (∼2.5 fold) within five years of diagnosis and poorer overall survival rate (4.2 vs. 6 years) compared to receptor positive breast cancer subtypes^3,4^. This poor outcome derives from the heterogeneous nature of the disease, coupled with the lack of highly recurrent and/or actionable biomarkers that are informative for therapy^5,6^. Furthermore, while some TNBC are initially chemosensitive, 23% of patients recur within 5 years from diagnosis and over 30% develop drug resistance tumours^7,8^. Therefore, there is an urgent need to improve our understanding of the molecular basis for TNBC development and progression to discover effective therapeutic targets and their companion test to improve the outcome in patients.

Metabolic adaptation is inherent to tumorigenesis to meet the increased requirements for bioenergetic, biosynthetic, and detoxification demands of malignant cells^9^. An increased aerobic glycolysis rate is a common metabolic feature in many cancer cells, and has been under extensive investigation as a therapeutic focus in cancer^10,11^. Among breast cancers, TNBC cells have an elevated glycolytic gene signature and concomitant lower oxidative phosphorylation signature compared to other breast cancer subtypes such as the hormone-positive luminal breast cancer^12^. High expression of glucose transporter 1 (GLUT1), a key rate-limiting factor for glucose uptake, is significantly elevated in basal-like breast cancer subtype^13^ (the most common type of TNBC^14^). This suggests a key role for GLUT1 in regulating TNBC cell metabolism. Targeting GLUT1 with small molecules, such as STF-31, WZB-117 and BAY-876, has been investigated in various types of cancers with promising results^15–18^. GLUT1 inhibition, either by a short hairpin RNA (shRNA) or WZB-117 inhibitor treatment showed anti-proliferation effects in MDA-MB-231 and HS 578T TNBC cell lines, again supporting GLUT1 as a possible target in TNBC^19,20^. Hence, a more systematic investigation and better understanding of the mechanisms regulating GLUT1 dependency is urgently needed to assess the benefits of pharmacological inhibition of GLUT1 for TNBC treatment in preclinical settings.

The inherent plasticity of cellular metabolism and the high degree of metabolic heterogeneity in TNBCs pose great challenges for metabolism targeting therapy^21^. Recent work suggests that heterogeneous metabolic dependencies within cancer cells underline the differential therapeutic vulnerabilities^22^. Therefore, in order for GLUT1 inhibition to be a successful strategy for TNBC therapy, the precise contexts in which this metabolic pathway is essential needs to be identified. In addition to tumor microenvironment, metabolic dependencies can be driven by genetic lesions, such as *myc* amplification and *Kras* mutation^23,24^. However, this oncogene driven cancer metabolism is incredibly complex and context-specific across cancer types^23,25–26,27^. Reliable biomarkers for predicting GLUT1 dependence and GLUT1 inhibition sensitivity are still lacking in TNBC.

In this study, we systematically assessed the vulnerability of a wide range of well-characterized TNBC cell lines to GLUT1 pharmacological inhibition. We then identified the molecular basis underlying GLUT1 dependencies, and validated our results in patient derived organoids and tumour explants. Finally, we identified RB1 protein levels as a predictive biomarker for GLUT1 sensitivity, which may potentially be used to stratify TNBC patients that would benefit from targeted GLUT1 therapy.

## Results

### Growth of a subset of TNBC relies on GLUT1 activity

To test the GLUT1 dependency of TNBC, we first investigated whether the expression level of *SLC2A1*, the gene encoding GLUT1, was increased in TNBC by interrogating two large independent publicly available clinical cohorts, the TCGA and METABRIC^28,29^. In both cohorts examined, *SLC2A1* mRNA expression is significantly elevated in basal-like subtype (corresponding to the most common subtype of TNBC^13^) compared to oestrogen receptor positive and HER2-amplified breast tumors (TCGA: p=3.33e-11; METABRIC: p=2.53e-8.) (Fig. 1a-b). Similarly, *SLC2A1* elevated mRNA levels were observed in a smaller, independent breast cancer patient derived xenograft (PDX) cohort from the Princess Margaret Cancer Centre (PM-PDXs) (Fig 1c. p=1.67e-2). PAM50 based breast cancer subtype classification across these datasets also revealed increased *SLC2A1* mRNA expression levels in the basal-like subtype over all other subtypes (Supplementary Fig. 1a-c)^30^.

**Fig. 1:**
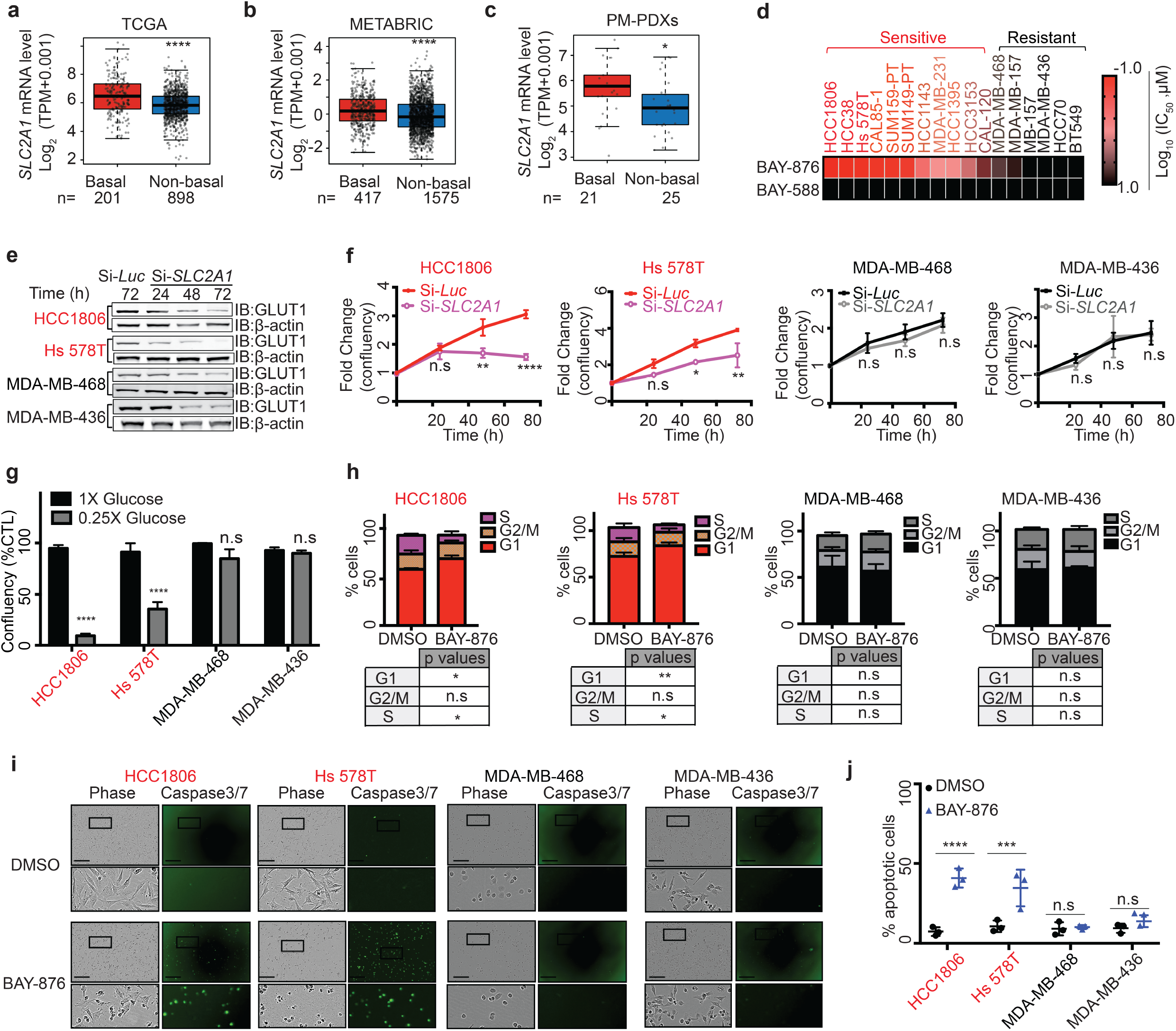
Growth of a subset of TNBC relies on GLUT1 activity. *SLC2A1* gene expression in the **(a)** TCGA Breast cancer datasets, **(b)** METABRIC Breast cancer datasets, and **(c)** Princess Margaret Hospital PDXs datasets (PM-PDXs). According to PAM50 classification, the cohorts were designated as basal and non-basal subtypes. Gene expression is reported as log2(TPM+0.001). The number of patients (n) per group is indicated. *P* values were determined using a Wilcoxon rank sum test. * p<0.05; ****p<0.0001. **(d)** Heatmap of mean IC_50_ values for the indicated 17 TNBC cell lines. Cells were treated with increasing dose of BAY-876 or its negative control BAY-588 for 5 days and the cell confluency was determined by Incucyte scanning. Data shown are mean ± s.d. of n = 4 independent biological replicates. A two-sided Student’s t-test was used to derive the p values. **(e)** Representative immunoblots showing the siRNA knockdown of GLUT1 in the BAY-876 sensitive lines (HCC1806 and Hs 578T) and BAY-876 resistant lines (MDA-MB-436 MDA-MB-468). **(f)** Normalized cell confluency of GLUT1 knockdown cells or siRNA luciferase control cells for the indicated time post siRNA transduction. Cell confluency are normalized to T=0 time point. Data shown are mean ± s.d. of n = 4 biological replicates. A two-way ANOVA was used to derive the p values. *p<0.05; **p<0.01; ****p<0.0001; n.s means not significant. **(g)** Cell growth of TNBC lines cultured in complete DMEM medium with or without glucose deprivation for 5 days. Data shown are mean ± s.d of n = 4 independent experiments. P values computed using a two-way ANOVA. ****p<0.0001. **(h)** Flow cytometry cell cycle analysis for indicated cells cultured with or without BAY-876 for 72h. Data shown are mean ± s.d. of n = 3 independent assays. A two-way ANOVA was used to derive the p values. *p<0.05; **p<0.01; ***p<0.001; n.s means not significant. **(i)** Representative images of caspase 3/7 staining. scale bar represents 300 μM. **(j)** Apoptotic cell counts of BAY-876 treated for 3 days by caspase 3/7 staining. Data shown are mean ± s.d. of n = 3 independent experiments. A two-way ANOVA was used to derive the p values. ***p<0.001; ****p<0.0001; n.s means not significant.

To further assess the function of GLUT1 as a target and the feasibility of GLUT1 inhibition as a therapy for TNBC patients, we treated a panel of 17 TNBC cell lines with the small molecule GLUT1 inhibitor BAY-876. Among the reported GLUT inhibitors, BAY-876, is the only inhibitor that is both highly potent and selective for GLUT1 over other glucose transporters (Supplementary Table. 1). Significant growth inhibitory effects were observed in 11 of the 17 TNBC cell lines (Fig. 1d and Supplementary Fig. 1d) based on a half maximal inhibitory concentration (IC_50_) value of less than 5 μM (range from 0.1 to 4.5 μM). The IC_50_ value for BAY-876 was greater than 10 μM for the six ‘resistant’ TNBC cell lines, the maximum dose used in the treatment. To complement the results from short-term treatments, we performed long-term (14 days) colony-forming assays to determine if the inhibitory effects of BAY-876 are sustained over time. At BAY-876 concentrations of 1 μM, proliferation of sensitive cell lines (HCC1806 and Hs 578T) was severely inhibited, whereas the resistant cell lines showed little effect (MDA-MB-436 and MDA-MB-468) (Supplementary Fig. 1e). These data confirm the heterogeneous response to GLUT1 pharmacologic inhibition across TNBC cell lines. We next evaluated the effect of siRNA-mediated silencing of GLUT1 on cell proliferation. Consistent with the results from pharmacological inhibition of GLUT1 with BAY-876, *SLC2A1* silencing reduced GLUT1 protein levels (Fig. 1e) and significantly impaired the growth of TNBC cell lines sensitive to BAY-876 (HCC1806 and Hs 578T) but had no impact on the growth of BAY-876 resistant TNBC cell lines (MDA-MB-436 and MDA-MB-468) (Fig. 1f). In agreement, partial deprivation of glucose from the culture media selectively impaired the growth of cell lines sensitive to BAY-876 treatment but had no significant effect on the BAY-876 resistant cell lines over 5 days (Fig. 1g).

We next characterized the mechanism of BAY-876 impaired growth in TNBC cell lines by quantifying the impact on cell cycle and apoptosis. The BAY-876 sensitive HCC1806 and Hs 578T cell lines demonstrated a modest but significant decrease in the S phase, with a concurrent increase in G1 phase following 72 hours treatment with 3 μM BAY-876 (Fig. 1h). In contrast, MDA-MB-436 and MDA-MB-468 cells showed no significant changes in cell cycle progression (Fig. 1h). Moreover, caspase 3/7 staining showed a significant increase in the number of apoptotic cells in BAY-876 sensitive compared to resistant cell lines upon GLUT1 inhibition (Fig. 1i-j). Taken together, these data showed that GLUT1 inhibition either by siRNA mediated GLUT1 silencing or by pharmacological inhibition using BAY-876 treatment, results in attenuated cell growth and proliferation, increased cell cycle arrest and increased cell apoptosis, which collectively contribute to growth suppression in a subset of TNBC cells.

### OXPHOS levels correlate with the response to GLUT1 inhibition

As our data indicated that BAY-876 treatment selectively impairs the growth of a subset of TNBC cell lines, we assessed the mechanism conferring this heterogeneous response to GLUT1 inhibition. Because glucose is the fuel for glycolytic cellular metabolism, we reasoned that sensitivity to GLUT1 inhibition may be connected to the basal metabolic state of each cell line. Bioenergetic profiling revealed that the basal glycolytic rate as reflected by the extracellular acidification rate (ECAR) and mitochondrial oxygen consumption rates (OCR) indicative of oxidative phosphorylation (OXPHOS), discriminates between BAY-876 sensitive versus and resistant TNBC cell lines (Fig. 2a). Whereas resistant cell lines exhibited slightly decreased ECAR (glycolytic rates), they display a 3-fold higher OCR (oxygen consumption rate) compared to sensitive cell lines at the basal level (in absence of BAY-876) (Fig. 2a). The ratio of OCR to ECAR (OCR/ECAR), indicative of higher reliance on OXPHOS, was significantly higher in resistant compared to sensitive TNBC cell lines (Fig. 2b)^31^. This observation indicates that BAY-876 resistant cells display higher levels of OXPHOS at the basal state compared to BAY-876 sensitive TNBC cell lines.

**Fig. 2:**
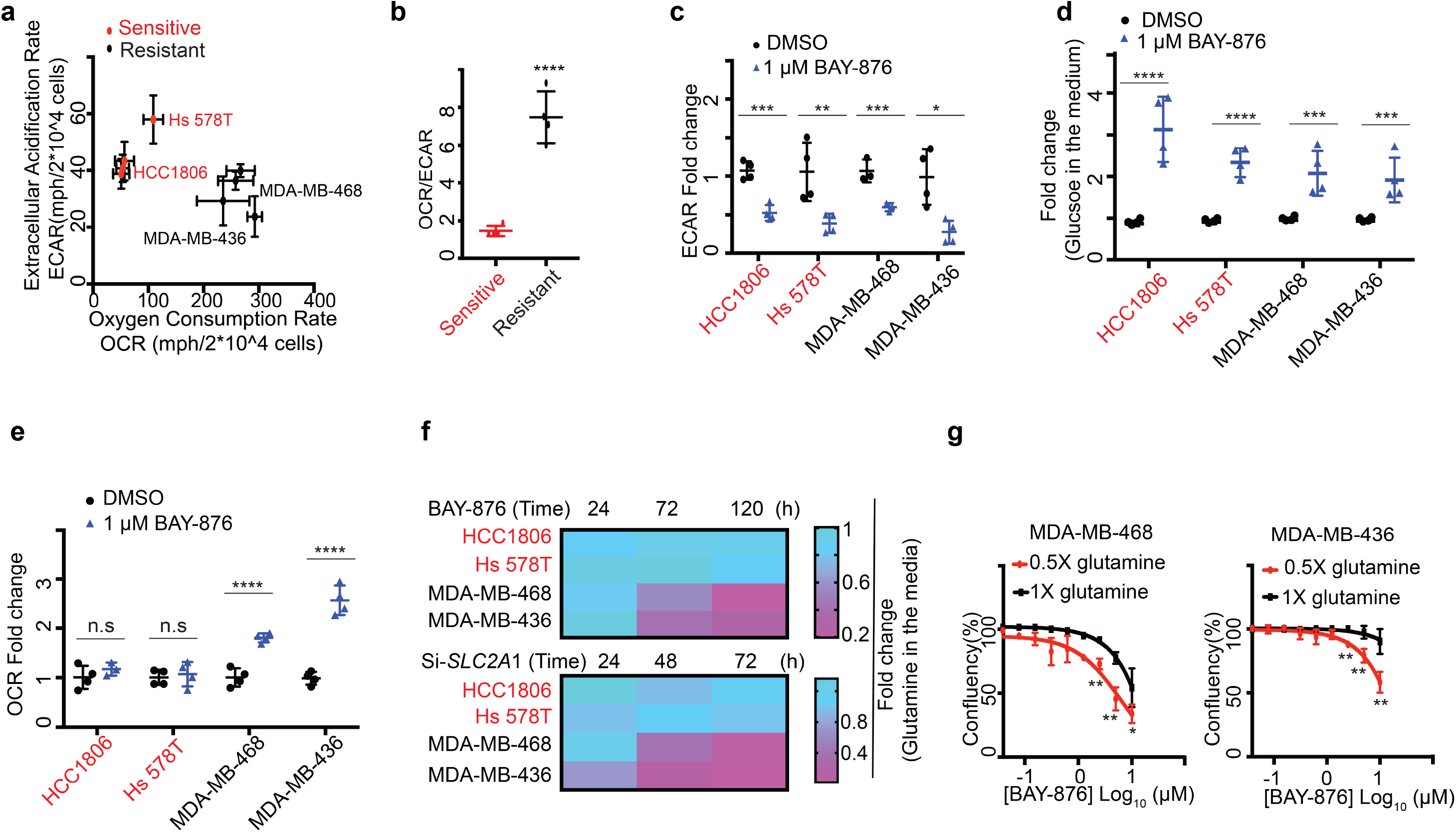
OXPHOS levels correlate with the response to GLUT1 inhibition. **(a)** OCR and ECAR were measured for each of BAY-876 sensitive lines (red) and BAY-876 resistant lines (black) using XF-96 analyzer. ECAR and OCR values were normalized to total cell numbers for each cell line. Plots show mean values from nine wells (from three experiments) and relative ECAR and OCR data were plotted simultaneously to reveal overall relative basal metabolic profiles for each cell model. **(b)** OCR and ECAR ratio were calculated for each cell line. P values were derived from a two-sided Student’s test; ****p<0.0001. **(c)** ECAR values (n=4; mean ± s.d.) measurement of cells with or without BAY-876 treatment for 5 days. P values computed using a two-way ANOVA. *p<0.05; **p<0.01; ***p<0.001. **(d)** Glucose uptake analysis using Bioprofile Flex analyzer (Nova Biomedical) were performed in cells following BAY-876 treatment for the indicated time. Data shown are mean ± s.d. of n = 4 independent assays. P values computed using a two-way ANOVA. ***p<0.001; ****p<0.0001; **(e)** A trace of OCR values (n=4; mean ± s.d.) from a mitochondrial stress test of cells with or without BAY-876 treatment for 5 days. P values computed using a two-way ANOVA. ****p<0.0001; n.s denotes not significant. **(f)** Glutamine uptake analysis using Bioprofile Flex analyzer (Nova Biomedical) were performed in cells following 1 μM BAY-876 treatment (n=3) or in cells transfected with 25nM siGLUT1 (bottom) (n=2). **(g)** Growth curves of MDA-MB-468 cells (left) and MDA-MB-436 cells (right) cultured in complete DMEM medium with or without glutamine deprivation treated with indicated nine doses of BAY-876 for 5 days. Data shown are mean ± s.d of n = 3 independent experiments. P values computed using a two-way ANOVA. **p<0.01; ***p<0.001; ****p<0.0001.

In addition to the basal metabolic bioenergetic profile, a subclass of breast cancer cells have also been reported to switch from aerobic glycolysis to OXPHOS under limiting glucose conditions, as observed in cervical cancer, glioma and pancreatic cancer cells^32–34^. This metabolic plasticity illustrates the interplay between glycolysis and OXPHOS, enabling the cells to adapt their bioenergetic profile to microenvironmental changes^35^. In agreement with BAY-876 inhibiting glucose uptake, the rate of glycolysis was significantly decreased upon GLUT1 inhibition as measured by decreased ECAR (Fig. 2c), glucose uptake (Fig. 2d) and lactate secretion (Supplementary Fig. 1g) in both BAY-876 sensitive and resistant TNBC cell lines. However, BAY-876 resistant cell lines display approximately double OCR upon GLUT1 inhibition, while no significant difference in OCR was observed in sensitive TNBC cell lines (Fig. 2e). This suggests that BAY-876 resistant cells can adopt an increased OXPHOS metabolic profile to compensate for decreased glucose uptake, thereby enabling continued cell growth and cell survival. Since glutamine is utilized as a major energy source to drive OXPHOS, we next tested the dependence of both sensitive and resistant TNBC cell lines on glutamine. BAY-876 resistant TNBC cell lines exhibited glutamine depletion in the media (indicative of increased glutamine uptake) either upon *SLC2A1* knockdown leading to depleted GLUT1 levels or BAY-876 treatment (Fig. 2f). In addition, removal of glutamine from the growth medium resulted in an increased sensitivity to BAY-876 in resistant TNBC cell lines, suggesting a strong dependence of resistant cells to glutamine-fueled OXPHOS to bypass growth suppression induced by GLUT1 inhibition (Fig. 2g). These results further support the ability of BAY-876 resistant TNBC cells to adapt their bioenergetic profile and metabolic requirements upon blocking GLUT1.

### RB1 protein level discriminates response to GLUT1 inhibition

GLUT1 is known to influence a wide variety of biological processes, however, it is still unclear how these underlie the GLUT1 dependency of cancer cells^36,37^. To address this key question, we first examined molecular and phenotypic features that correlated with sensitivity or resistance to BAY-876 using the IC_50_ values calculated from our 17 TNBC cell lines (Fig. 1d). The highly reproducible responses identified in both sensitive and resistant cell lines, indicate that BAY-876 treatment and GLUT1 inhibition are not universally cytotoxic. To identify the molecular mechanism(s) for cellular drug resistance, we considered several candidate pathways. Since BAY-876 efficiently decreased glucose uptake and glycolysis rates in highly resistant cell lines (Fig. 2cd), we ruled out drug-efflux pump or impaired drug metabolism mechanisms which could decrease the effective cellular concentration of BAY-876. Next, we examined the mRNA expression of *SLC2A1* across all 17 cell lines. The response to BAY-876 is not correlated with *SLC2A1* mRNA levels (Supplementary Fig. 2a). However, it is difficult to draw conclusions from this observation given that total *SLC2A1* expression does not necessarily correlate with its protein level^38^. We then performed a systematic global profiling of the published transcriptome data for our panel of TNBC cell lines^39^. We profiled differentially expressed genes between BAY-876 responders and nonresponders (Fig. 3a) and subjected these genes to gene set enrichment analysis (GSEA). Gene sets associated with the OXPHOS pathway stood out in the analysis as enriched in BAY-876 resistant versus sensitive TNBC cell lines (Fig. 3b and Supplementary Fig. 2b), suggesting that the elevated functional mitochondrial output of resistant cells as shown in Fig. 2 is due to the increased OXPHOS gene expression signature.

**Fig. 3:**
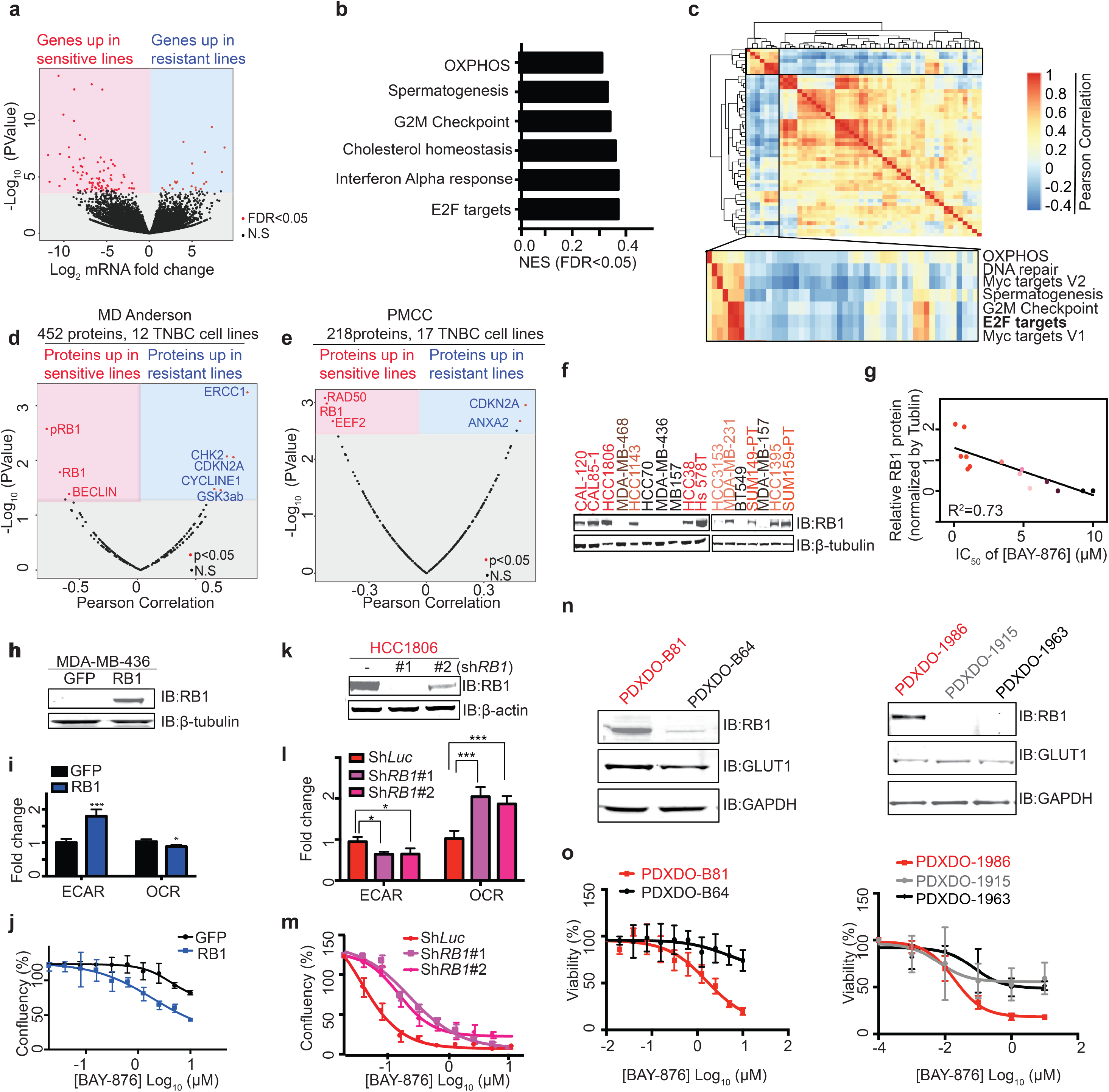
RB1 protein level discriminates response to GLUT1 inhibition. **(a)** Volcano plot of log2 fold change for all genes significantly upregulated (red in the left panel) or downregulated (blue in right panel) in sensitive compared to resistant lines. **(b)** Top enriched pathways in BAY-876 resistant lines compared to BAY-876 sensitive lines based on GSEA on RNA-sequencing data generated from UHN cohort. **(c)** Heatmap of pathways correlated with OXPHOS revealed by GSEA analysis of TCGA RNA-sequencing data. Color grading corresponds to the positive (red) and negative (blue) Pearson’s correlation coefficients. Rows and columns were hierarchically clustered by Euclidean Distance with complete linkage. Clusters significantly related with OXPHOS are zoomed at the bottom. **(d)** Pearson’s correlation coefficients between Log2 normalized protein expression data and response of BAY-876 showing significantly associated proteins (red) with BAY-876 sensitive and resistant lines based on the dataset from MD-Anderson Cancer Center including 12 TNBC cell lines and 452 proteins. **(e)** Pearson’s correlation coefficients between Log2 normalized protein expression data and response of BAY-876 showing significantly associated proteins (red) with BAY-876 responders and non-responders based on the dataset from Princess Margaret Cancer Center (PMCC) including 17 TNBC cell lines and 218 proteins. **(f)** Representative western blot showing the variable RB1 expression levels in a panel of 17 TNBC lines. β-actin was used a loading control for normalization. **(g)** Relative RB1 protein levels in TNBC, quantified by densitometry from three independent experiments were plotted with the IC_50_ of BAY-876. **(h)** Representative immunoblot showing MDA-MB-436 cells expressing RB1 or GFP control proteins. β-actin was used as a loading control. **(i)** ECAR and OCR values were measured for MDA-MB-436 cells expressing RB1 or GFP control. Data shown are mean ± s.d of n = 4 independent biological replicates. P values computed using a two-sided Student’s t test. *p<0.05; ***p<0.001. **(j)** Growth curves of MDA-MB-436 cells expressing RB1 or GFP control in the presence of indicated concentrations BAY-876 treatment for 5 days. Data shown are mean ± s.d of n = 4 independent biological replicates. **(k)** Representative western blot showing HCC1806 cells transfected with two independent shRNAs targeting RB1 or shRNA targeting control luciferase. β-actin was used as a loading control. **(l)** ECAR and OCR values were measured for HCC1806 cells transfected with two independent shRNAs targeting RB1 or shRNA targeting control luciferase. Data shown are mean ± s.d of n = 4 independent experiments. P values computed using a two-way ANOVA. *p<0.05; ***p<0.001. **(m)** Growth curves of HCC1806 cells with control knockdown or RB1 knockdown in the presence of indicated concentrations BAY-876 treatment for 5 days. Data shown are mean ± s.d of n = 4 independent experiments. **(n)** Western blot showing the variable RB1 expression levels in a panel of five TNBC patient-derived organoids. GAPDH was used as a loading control. **(o)** Cell viability assays five days after administration of DMSO control or the indicated concentrations doses of BAY-876 across five independent patient-derived organoids. Data shown are mean ± s.d of n = 3 independent biological replicates.

GSEA analysis also revealed that the most significantly enriched pathway in resistant versus sensitive cell lines is the E2F Targets pathway (Fig. 3b and Supplementary Fig. 2c), suggesting that elevated expression of genes involved in the E2F Targets pathway also correlates with resistance to GLUT1 inhibition. To further confirm this association in patient samples, we took advantage of the transcriptome data of patient TNBC tumors from TCGA data cohorts. Based on our observation of strong correlation between elevated OXPHOS metabolism and resistance to GLUT1 inhibition in TNBC cell lines, we used increased gene expression patterns of OXPHOS to discriminate BAY-876 putative-resistant samples versus putative-sensitive samples in the TCGA cohorts. Calculating the Pearson correlation coefficients between the OXPHOS pathway with other gene expression patterns revealed a strong clustering with the E2F Targets pathway, suggesting that high expression of genes involved in the E2F Targets pathway correlates with expression of OXPHOS related genes in primary TNBC tumour samples (Fig. 3c and Supplementary Fig. 2d).

We next sought to identify a protein signature associated with the above gene expression and metabolic differences and that could predict the relative responsiveness of TNBC cells to BAY-876. Using proteomics datasets from University of Texas MD Anderson Cancer Center for 12 of our TNBC cell lines^40^, we identified differential protein levels for a total of 8 proteins which showed a significant correlation with BAY-876 response (Fig. 3d). Most proteins (3 out of 5) enriched in resistant cell lines are components of the E2F Targets pathway, namely cyclin-dependent kinase inhibitor 2A (CDKN2A), cyclin E1 (CCNE1) and checkpoint kinase 2 (CHK2). Among the three proteins whose levels were increased in BAY-876 sensitive lines, the top hit was retinoblastoma tumor suppressor (RB1) protein that functions primarily as an upstream transcription factor attenuating expression levels of known E2F targets^41,42^. We further confirmed the association of RB1 protein levels with BAY-876 sensitivity in an independent proteomics datasets from the Princess Margaret Cancer Centre^39^ (Fig. 3e). We also assessed RB1 protein levels across our 17 TNBC cell lines by immunoblotting (Fig. 3f). This further confirmed the significant correlation (R^2^=0.73) between RB1 protein levels and BAY-876 sensitivity (Fig. 3g). These results suggest that elevated RB1 protein levels underly sensitivity to BAY-876 treatment, while low RB1 protein levels associates with resistance to BAY-876 in TNBC. To test this concept, we overexpressed RB1 in two BAY-876 resistant cell lines: MDA-MB-436 and MDA-MB-468 (Fig. 3h and Supplementary Fig. 3a). In both cell lines, RB1 overexpression caused an increase in the ECAR/OCR ratio and markedly sensitized cells to BAY-876 treatment (Fig. 3i-j and Supplementary Fig. 3b). Conversely, RB1 knockdown in BAY-876 sensitive TNBC cell lines induced higher OCR/ECAR ratios and rendered the cells refractory to the anti-proliferative effects of BAY-876 (Fig. 3k-m and Supplementary Fig. 3c-d). Collectively, our results suggest that RB1 protein levels can serve as a biomarker of response to BAY-876 treatment in TNBC cell lines.

### Pharmacological inhibition of GLUT1 impedes TNBC cancer growth in patient-derived models

To better address the clinical relevance of this hypothesis, we tested the correlation of RB1 protein level and BAY-876 sensitivity across a panel of TNBC patient derived samples. Patient-Derived Xenograft (PDX)-Derived Organoids (PDXDOs), which are thought to better recapitulate features of breast histology and epithelial heterogeneity, and therefore serve as better tools to assess drug responses for cancer therapy^43^. Thus, PDXDOs from five different TNBC patients were cultured, of which three cases are RB1-low, and two expressed RB1 based on immunoblots (Fig. 3n). In agreement with our observations in TNBC cell lines, higher RB1 expression was predictive of sensitivity to BAY-876 in PDXDOs (Fig. 3o). Some PDXDOs (PDXDO-B64, PDXDO-1915 and PDXDO-1963) with low-RB1 levels showed a weakened partial response to BAY-876 treatment. We postulated that this could be due to heterogeneity in RB1 expression across sub-populations of cells found in each patient-derived tumour culture. In agreement, Immunostaining for RB1 protein indicates that although most of the cells are RB1 negative in PDXDO-1915 (85.9% cells are RB1 negative) and PDXDO-B64 (65.7% cells are RB1 negative), some cells showed a strong RB1 staining signal confirming a mixed population with differential RB1-related response to BAY-876 (Supplementary Fig. 3e-f).

We further tested the efficacy of GLUT1 inhibition and its association with RB1 protein expression in PDX derived tumor *ex-vivo* explant models (PDXDEs) (Fig. 4a). PDXDE models have been shown to accurately predict patient-specific responses to multiple drugs and to mimic *in vivo* results in breast cancer^44,45^. We first established the appropriate culture conditions to maintain TNBC PDXDEs. As illustrated by hematoxylin and eosin (H&E) staining, tissue architecture and morphology of PDXDEs cultured for up to 48 hours on gelatin sponges were consistent with the original (T=0h) tumor tissue (Fig. 4b and Supplementary Fig. 4a). Tumor cells are present in the surrounding stroma, demonstrating maintenance of the PDX architecture (Supplementary Fig. 4). The proliferative capacity of explants was assessed using the immunohistochemistry marker ki67, with a change of 25% or more considered to be a significant response^46^. No significant change in the number of ki67 positive cell nuclei was observed between T=0h and 48 hours of culture (T=48h) for matched tissues (Fig. 4c). Furthermore, as shown by immunohistochemistry staining for the apoptotic marker cleaved caspase-3 (CIC3), no significant differences in the proportion of apoptotic cells was observed in PDXDEs cultured over a 48 hours period (Fig. 4d). Altogether, our results demonstrate that PDXDEs are viable over the experimental period of 48 hours.

**Fig. 4:**
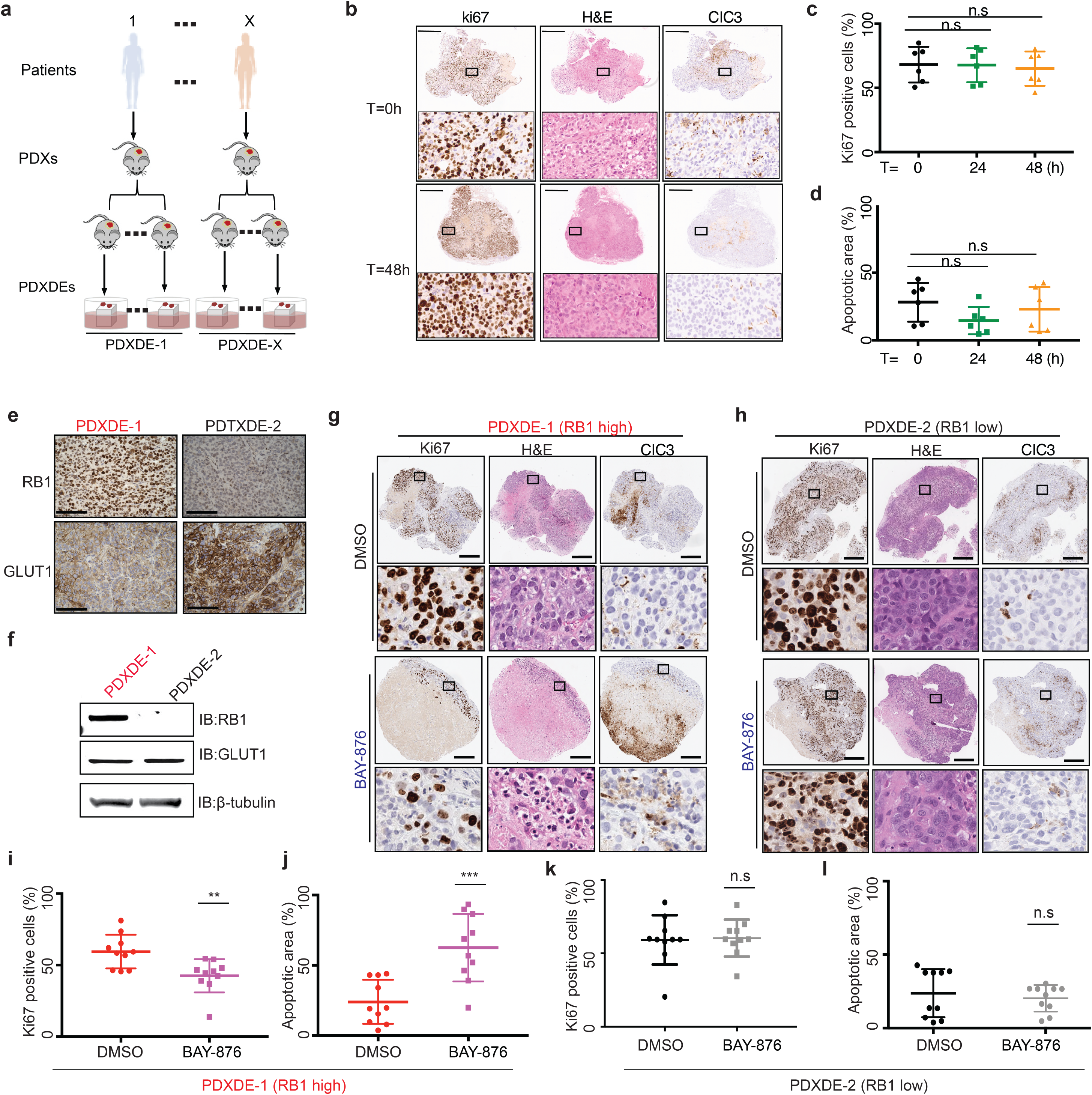
RB1 levels dictates BAY-876 sensitivity in patient derived explants. **(a)** Schematic of pre-clinical PDXDE trial. PDXDEs established from TNBC patients were evaluated for response to BAY-876 treatment. **(b)** Representative images of explants during *ex vivo* culture time range assessed by proliferative index ki67 staining, H&E staining and cleaved caspase 3 (CIC3) staining. Scale bars represent 500 μm in the representative IHC staining images. Indicated area is zoomed in 5X at the bottom. **(c)** *Ex-vivo* culture for 48 hours did not significantly change the cell proliferation of explants assessed by Ki-67 staining. Data shown are mean ± s.d of n = 6 independent experiments from three biological replicates. **(d)** *Ex-vivo* culture for 48 hours did not significantly change the apoptosis of explants. Data shown are mean ± s.d of n = 6 independent experiments from three biological replicates. **(e)** Representative IHC staining images of GLUT1 and RB1 for PDXDE-1, PDXDE-2. **(f)** Representative immunoblotting showing RB1 and GLUT1 expression levels in PDXDE-1 and PDXDE-2. **(g)** Representative IHC staining images including ki67 staining, H&E staining and CIC3 staining of RB1-positive PDXDE-1. Scale bars represent 500 μm. Indicated area is zoomed in 10X at the bottom. **(h)** Representative IHC staining images including ki67 staining, H&E staining and CIC3 staining of RB1-low PDXDE-2. Scale bars represent 500 μm. Indicated area is zoomed in 10X at the bottom. **(i)** BAY-876 treatment resulted in regression of PDXDE-1 growth assessed by ki67 staining. Data shown are mean ± s.d of n = 10 independent experiments from three biological replicates. P values computed using a two-sided Student’s test. ****p<0.0001. **(j)** BAY-876 treatment resulted in increased apoptosis of PDXDE-1 assessed by CIC3 staining. Data shown are mean ± s.d of n = 10 independent experiments from three biological replicates. P values computed using a two-sided Student’s test. ***p<0.001. **(k)** BAY-876 treatment did not result in significant change of PDXDE-2 growth assessed by ki67 staining. Data shown are mean ± s.d of n = 11 independent experiments from three biological replicates. P values computed using a two-sided Student’s test. **(l)** BAY-876 treatment did not lead to significant changes of apoptosis of PDXDE-2 assessed by CIC3 staining. Data shown are mean ± s.d of n = 10 independent experiments from three biological replicates. P values computed a twosided Student’s test.

Next, we quantified the RB1 protein level by RB1 immunohistochemistry staining in PDXDEs from six different patients (Supplementary Fig. 5a)^47,48^. Each PDXDE case was assigned a score of 0 (<10% of cells positively stained), 1 (if >10% and <50% of cells positively stained) or 2 (>50% of cells positively stained), representing RB1-negative, intermediate or positive PDXDEs respectively, (Supplementary Fig. 5a). The RB1-positive (PDXDE-1) and RB1 negative explants (PDXDE-2) were then used to evaluate the efficacy of GLUT1 inhibition. Each of these PDXDEs expressed GLUT1 at similar levels (Supplementary Fig. 5b). RB1 and GLUT1 expression levels in both PDXDEs were confirmed by immunoblotting (Fig. 4e). Consistent with our cell line and PDXDO results, BAY-876 (3 μM) treatment abrogated the proliferation in 70% (7/10) PDXDE-1 RB1-positive explants within 48h as measured by ki67 staining when compared to vehicle-treated PDXDEs (Fig. 4g, Supplementary Fig. 6). Although intertumoral heterogeneity to BAY-876 response was observed across these explants, PDXDE-1 explants showed a significant overall reduction in ki-67 proliferation (p<0.01) upon BAY-876 treatment (Fig. 4i). Moreover, cleaved caspase-3 staining revealed a significant increase in apoptosis in BAY-876 treated PDXDE-1 (Fig. 4j and Supplementary Fig. 6). Conversely, no significant changes in cell proliferation or apoptotic markers were observed following BAY-876 treatment in the RB1-negative PDXDE-2 explants (Fig. 4h, 4k-l and Supplementary Fig. 7). Collectively, our findings across a range of pre-clinical PDX derived models provide a strong rationale for selecting GLUT1 inhibition as a therapeutic strategy on the basis of RB1 protein expression level and supports the further evaluation of strategies to inhibit GLUT1 and/or the RB1/E2F axis in TNBC patients.

## Discussion

In this study, we identify a dependency of a subset of TNBCs on GLUT1 function, and relate this dependency to the distinct basal cellular bioenergetic profile. Our findings suggest a mechanistic basis by which cells with higher glycolysis/OXPHOS rate are susceptible to GLUT1 inhibition. These results have important implications for the design of therapeutic strategies for TNBC, given the known heterogeneity within TNBCs, both genetically and metabolically^49^. This is also supported by the recent observations of metabolic heterogeneity across diverse cancer cell lines at both the unperturbed and the perturbed states^22^. The latter study suggests that heterogeneous metabolic dependencies across cancer cell lines underly differential therapeutic vulnerabilities associated with specific cancer genotypes^22^. Direct targeting of metabolic states in TNBC has shown promising results, for example, inhibitors that target glutathione biosynthesis, folate receptor, fatty acid oxidation as well as glutamine metabolism were shown to suppress tumor growth in TNBC^26,27,50,51^. However, successful clinical translation of these results is hampered by issues such as metabolic plasticity ^21^.

Metabolic plasticity need not be inherent but may be adaptive, based on the stage of tumor progression, tumor microenvironment such as nutrient availability, and the type of treatment administered^52,53^. It is likely that in many cases in which glycolysis is inhibited, cells will respond by increasing other alternative metabolic pathways^54,55^. Supporting this, our results demonstrated that GLUT1 inhibition triggered metabolic reprogramming toward OXPHOS in BAY-876 resistant TNBC cell lines. This metabolic adaptation in conjunction with the use of alternative nutrients such as glutamine overcomes the inhibition of glucose metabolism in our BAY-876 resistant TNBC cell lines. Thus, glutamine deprivation sensitizes the resistant cells to BAY-876 inhibition. This synergism between glycolysis inhibitors combined with OXPHOS inhibition suggests a strong rationale using two or more drugs targeting different metabolic pathways to achieve superior therapeutic benefits for TNBC treatment^56,57^.

Many relationships between cancer metabolism and genotypes have been established in which genomic alterations are used to identify metabolic differences^58,59^. Mutations that activate oncogenes or inactivate tumor suppressors can significantly affect activities of metabolic enzymes and have a key role in aerobic glycolysis of cancer^23,24,60-62^. Phosphatidylinositol 3’-kinase (PI3K)^63^, phosphatase and tensin homolog (PTEN)^64^, Myc^65,66^ and p53^67^ can all impact cellular glucose metabolism. Here, we discovered the significant correlation between RB1-E2F Targets pathway and OXPHOS in both cell lines and TNBC primary tumors. TNBC cells that lack RB1 protein have increased expression of OXPHOS genes and accordingly increase mitochondrial respiration, ultimately leading to resistance to GLUT1 inhibition. This metabolic role of RB1 in TNBC is under appreciated, but consistent with recent reports suggesting that RB1 loss stimulates mitochondrial function rather than anaerobic metabolism by activating E2F targets^68,69^. These connections are especially interesting given the rapidly growing evidence that RB1 is needed for cells to maintain a normal metabolic balance, and that the loss of RB1 leads to reprogramming of specific pathways^70–72^. Metabolic adaptations are thought to enhance the ability of cancer cells to sustain the metabolic intermediates for cell survival during multidrug resistance^73,74^. Our results outline a clinically feasible scenario to use RB1 as a biomarker to stratify the TNBC patients for targeted therapy. Since RB1 is controlled by both genetic and epigenetic mechanisms, the presence of genetic mutations in RB1 fails to predict RB1 protein level^75,76^. Thus, we suggest using immunohistochemical assessment of RB1 protein in patient tumors to select patients suitable for targeted therapy in TNBC.

In summary, we have discovered a RB1 protein-dependent metabolic addiction to GLUT1 function in a subset of TNBCs, identifying BAY-876 as an effective agent to block growth in patient derived models, including explants that express RB1 protein.

## Methods

### Cell culture

Human breast cancer cell lines were obtained from American Type Culture Collection (ATCC, Teddington, UK). All cells were routinely cultured in RPMI 1640 (Life Technologies 11965), or Dulbecco’s modified Eagle’s medium (Gibco) supplemented with 10% FBS recommended by suppliers. The cell lines were authenticated by short-tandem-repeat (STR) analysis and matched to the German Collection of Microorganisms and Cell Cultures (DSMZ) database, and they were used for no more than 25 passages after STR typing. Mycoplasma tests were routinely performed using MycoAlert Mycoplasma Detection Kit (Lonza, Basel, Switzerland).

### SLC2A1 mRNA expression analysis

TCGA mRNA expression was downloaded from the Xena browser^77^. METABRIC mRNA expression was downloaded using MetaGx R package. Genefu R package^78^ was used to classify all samples into PAM50 molecular subtypes. Wilcoxon rank sum test was used to measure the significance of difference between TNBC vs non-TNBC samples and also between the different specific subtypes.

### TCGA data analysis

Breast Adenocarcinoma TCGA data was retrieved from Firehose using the GSVA R package^79^. We identified enrichment of Hallmark gene sets^80^ for each TNBC tumor sample in TCGA using their RNAseq profiles. The enrichment was conducted using ssgsea method as part of GSVA R package^79^. The correlation between the Hallmarks were then calculated using Spearman’s rank correlation.

### Proliferation and colony-formation assays

TNBC control and gene knocked down cells (500/well) were seeded in 384 well plate and transferred to Incucyte ZOOM analysis system (Sartorius) that was maintained at 37 °C. Growth profile was monitored by 10 X objective every 6 h using Incucyte software 2016 A with an integrated confluence algorithm until 72h. Standard mode per well was used to collect images in phase-contrast mode and averaged to provide a representative statistical measure of the well confluence. For colony-formation assays, 500–1,000 cells were seeded in 6-well plates. At the indicated time point (usually 10–14 days), cells were fixed with 80% methanol and stained with crystal violet solution overnight. All experiments were performed in triplicate.

### Small interfering RNA (siRNA)-mediated gene knockdown

siGENOME siRNA targeting *SLC2A1* (L-007509-02) and a non-targeting siRNA pool (D-001206-14) control, were purchased from Dharmacon (Thermo Scientific, Hemel Hempstead, UK. Cells were transfected with 25 nmol/L siRNA using Lipofectamine RNAiMax transfection reagent (Invitrogen, Life Technologies, Paisley, UK), following the manufacturer’s instructions.

### Western blot analysis

Cells were lysed directly in 1× lysis buffer (50 mmol/L Tris-HCl pH 6.8, 2% SDS, 10% glycerol, 2.5% β-mercaptoethanol and 0.1% bromophenol blue). Alternatively, cells were lysed by scraping them into a pH 7.4 lysis buffer containing 1% NP-40 (Sigma-Aldrich, Gillingham, UK), 50 mmol/L Tris, 10% glycerol, 0.02% NaN3, 150 mmol/L NaCl, and a cocktail of phosphatase and protease inhibitors (Sigma-Aldrich, Gillingham, UK. Snap-frozen tumor tissues were suspended in a pH 7.4 lysis buffer containing 50 mmol/L Tris base, 150 mmol/L NaCl, 2% TritonX-100, 1% SDS, 10 mmol/L EDTA, and a cocktail of phosphatase and protease inhibitors (Sigma-Aldrich, Gillingham, UK). Tissue destruction was done with the bullet blender homogenizer (Next Advance, New York, USA). 20–100 μg of proteins were separated in reducing conditions (2.5% β-mercaptoethanol) by SDS–PAGE (SDS–polyacrylamide gel electrophoresis) and transferred to nitrocellulose membranes (Bio-Rad, Hemel Hempstead, UK) for further processing, following standard western blotting procedures.

Primary antibodies used in this study were: anti-GLUT1 rabbit monoclonal (EPR3915) (ab115730) (1:1000), anti-RB1 rabbit monoclonal (EPR) (ab181616) (1:2000), anti-beta actin (ab16039) (1:5000). The secondary antibodies are goat-anti rabbit (IR800 conjugated, LiCor no. 926-32211) and donkey anti-mouse (IR 680, LiCor no. 926-68072) antibodies (1:5000). Odyssey Licor system were used to scan membranes and ImageJ was used to quantify western blotting results by densitometry

### Cell cycle assay

Exponentially growing cells in six-well plates were treated with 3 μM BAY-876 or DMSO for 24h before cell cycle analysis using allophycocyanin (APC) BrdU Flow kit (BD Pharmingen). Briefly, cells were incubated with 10 μM BrdU for 6 h before fixation, permeabilization, and staining with APClabeled anti-BrdU antibody and 7-aminoactinomycin D (7-AAD) according to the manufacturer’s instructions. Cells were then analyzed using a BD FACScan flow cytometer and the percentage of live cells in each cell cycle stage was determined using FlowJo software (version 9.3.1). Experiments were performed in duplicate and the percentage of cells in each stage was compared between the treated and untreated samples for each cell line using a two-tailed t test. P values ≤0.05 were considered significant.

### Cell apoptosis assay

For cell death assays TNBC cells (3000/ well) were seeded in 96 well plate and incubated for 24h at 37^0^C. The cells were then treated either with DMSO or BAY-876 for 5 days. Post treatment, Caspase-3/7 green apoptosis assay reagent (Sartorius #4440) was added to the cells and transferred to Incucyte^®^ ZOOM 2FLR system and analyzed using 2016 an integrated software.

### Mitochondrial respiration and glycolysis rate measurements

Bio-energetic studies (OCR: Oxygen Consumption Rate and ECAR: ExtraCellular Acidification Rate) were measured using a Seahorse XF^e^96 Extracellular Flux Bioanalyzer (Agilent). TNBC cells (10^4^) were seeded and cultured for 24h. The medium was then replaced with DMEM (25 mM glucose, 2 mM glutamine, no sodium bicarbonate) pH ∼7.4 and incubated for 1h at 37^0^C in a CO2-free incubator. For the mitochondrial stress test (Seahorse 101706-100), oligomycin, trifluoromethoxy carbonylcyanide phenylhydrazone (FCCP), and a mixture of antimycin and rotenone were injected to final concentrations of 2 μM, 0.5 μM and 4 μM, respectively. For the glycolysis stress test (Seahorse 102194-100), glucose, oligomycin and 2-deoxyglucose were injected to final concentrations of 10 mM, 2 μM and 100 mM, respectively. OCR and ECAR were normalized to cell number as determined by CyQUANT NF Cell proliferation assay kit (ThermoFisher Scientific, C7026). OCR and ECAR values were used to compute basal respiration, spare capacity, proton leak, ATP production, glycolysis and glycolytic capacity. Calculations from two independent experiments were performed using ExcelMacro Report Generator Version 3.0.3 provided by Seahorse Biosciences and two-sided Student’s *t*-test computed using GraphPad.

### Measurement of total cellular ROS

A dichlorofluorescin diacetate (DCFDA) cellular ROS detection assay (abcam, ab113851) was used to measure total ROS activity within the cells. A total of 2.5 × 10^4^ cells per well were seeded in a 96-well plate and allowed to attach overnight. The cells were then stained with 25 μM DCFDA for 45 min at 37 °C. After staining, the cells were washed and measured using a microplate reader with fluorescence (Ex/Em=485/535 nm).

### Metabolite measurements

Glucose, Lactate and Glutamine were quantified simultaneously using Bioprofile Flex analyzer (Nova Biomedical). Briefly, 300 μL of the cell/debris free culture medium was used to measure glucose/glutamine consumption and lactate production. The values were normalized to the cell number.

### Gene set enrichment analysis

Transcriptome data available for our tested 17 cell lines were processed using Kallisto pipeline (Marcotte et al. 2016, 10.1038/nbt.3519). Drug activity were extracted from growth curves produced by IncuCyte assay. IncucyteDRC R package was used to process the growth curves and obtain concentration and viability normalized to control near confluence point^81^ and PharmacoGx R package was then used to obtain drug activity measures such IC_50_ and Area-above-the-dose-response curve (AAC)^82^. Genes were ranked based on the Pearson correlation coefficients between the measured drug activity (IC_50_) and individual gene expression levels over all 17 cell lines. Hallmarks gene sets were downloaded from MsigDB^80^ and piano R package was used to produce the GSEA results. Pathways enrichment plots were generated using fgsea R package^83^.

### Proteomics analysis

MD Anderson protein expression data was downloaded from^40^. Princess Margaret Cancer Centre protein expression data was downloaded from^39^. Pearson correlation coefficients between the measured drug activity (IC_50_) and individual protein expression levels over all samples.

### Inducible ectopic expression of RB1

eGFP as control or RB1 were cloned into the Rc/CMV vector (Addgene plasmid 1763). Electroporesis-based transfection protocol (Lonza, previously known as Amaxa; http://www.lonzabio.com/cell-biology/transfection/) were used for transfection of MDA-MB-436 and MDA-MB-468 cells. 2–3 million cells were transfected with 3 micrograms of plasmid DNA vector. The transfection efficiency of this procedure TNBC cells amounted to 60%-80% and resulted in the transient expression of the transfected gene lasting up to 10–12 days.

### Lentiviral mRNA targets

Two independent shRNA vectors targeting RB1 were obtained from Addgene (Addgene ID: 25640 and 25641). Lentivirus was produced using standard virus production methods by co-transfecting target and packaging plasmids into HEK293T cells. Cell lines were then transduced with 0.45 μM filtered and ultracentrifuge-concentrated viral particles with Polybrene (8 μg ml–1). After 16 h of transduction, the media was changed for fresh regular growth media, and 48 h later selection started using puromycin (0.2–0.6 μg ml^−1^). After selection was complete in 72 h, cells were termed stably transduced.

### Immunohistochemistry staining

Paraffin sections at 4um thickness were dried at 60°C oven for 2 hours before staining. The immunohistochemistry (IHC) was performed according to the manufacturer’s guidelines using BenchMark XT-an automated slide stainer (Ventana Medical System). Glut1 (Roche #06419178001) IHC was done with mild antigen retrieval (CC1, pH8.0, #950-124), 32 min antibody incubation and Ventana iVeiw DAB Detection Kit (#760-091). The dilution for RB (BD #554136) was 1:1,600 with 64min antigen retrieval, 32min antibody incubation and Optiview Detection Kit (#860-099). Antibody dilution for Ki67 (Dako M7240, clone MIB1) was 1:100 with standard antigen retrieval, 60min antibody incubation and Ultraview Detection kit (#760-500). Dilution for cleaved caspase 3 (CST #9661) was 1:500 with standard antigen retrieval, 32min antibody incubation and iView Detection Kit. The slides were counterstained with Harris hematoxylin, dehydrated in graded alcohol, cleared in xylene and coverslipped in Permount.

### Image capture and quantification of immunostaining

Stained slides were subjected to whole slide imaging using an Aperio ScanScope AT2 at 20x magnification. Digital images were loaded into Definiens TissueStudio 4.3 software (Definiens Inc., Munich Germany), and individual tissue slices were identified away from slide background. A machine learning classifier was trained to identify viable tumor tissue from stroma, necrosis, and artifact, with manual quality correction to re-classify any mislabeled regions. Stain separation was used to isolate hematoxylin counterstain from DAB antibody-specific stain, and the hematoxylin channel was subjected to computer-vision based segmentation algorithm to identify individual nuclei, with a watershed step to break apart closely-packed nuclei. Cell simulation grew a small region around each nucleus to simulate a cytoplasm, and the resulting cell objects were classified into “negative”, “low” “medium’ and “high’ DAB intensity thresholds, based on comparison to clear positive and negative controls. Percell statistics were reported for each tissue region within each slide, and used to calculate the proportion of positive cells for each region of interest.

### Generation and maintenance of PDXDO-1915, PDXDO-1963 and PDXDO-1986

PDX were generated in NSG mice as above from TNBC biopsies and used under an REB-approved research protocol (UHN:15-9481). PDX tumors were excised from mice at passage 2, minced and digested in Advanced DMEM/F12 (ThermoFisher) containing 1X GlutaMAX, 10mM HEPES, 1X antibiotic-antimycotic and 500ug/mL Liberase TH (Sigma-Aldrich) in Miltenyi MACS C tubes. Digestion was performed using the gentle MACS Octo Dissociator (Miltenyi). Digested tissue was processed and plated in reduced growth factor Basement Membrane Extract (BME) Type 2 (Cultrex) in media as described by Sachs et al. (2018). The organoids were confirmed to be of human origin and devoid of mouse cells by flow cytometry. Organoids were passaged approximately every 3 weeks; they were dissociated to single cells with TrypLE Express (ThermoFisher) and by manual pipetting, passaged at a ratio of 1:12 and re-embedded in BME. For cell viability assays, organoids were dissociated and plated at 3,000 cells/well in 384-well plates pre-coated with 8mL BME. Cells were grown for 4 days; then, were treated with the indicated concentrations of BAY-876 for 5 days. Cell viability was assessed using Cell Titer Glo 3D (Promega).

### Generation and maintenance of PDXDO-B81 and PDXDO-B64

PDX tumors were excised from mice, minced and digested in Advanced DMEM/F12 (ThermoFisher) containing 1X GlutaMAX, 10mM HEPES, 1X antibiotic-antimycotic and 500ug/mL Liberase TH (Sigma-Aldrich) in Miltenyi MACS C tubes. Digestion was performed using the gentle MACS Octo Dissociator (Miltenyi). Digested tissue was processed and plated in reduced growth factor Basement Membrane Extract (BME) Type 2 (Cultrex) in media as described by Sachs et al. (2018). The organoids were confirmed to be of human origin and devoid of mouse cells by flow cytometry. Organoids were passaged approximately every 3 weeks; they were dissociated to single cells with TrypLE Express (ThermoFisher) and by manual pipetting, passaged at a ratio of 1:12 and reembedded in BME.

### *Ex vivo* tissue explant preparation and culture

When the PDXs (preferably first generation PDX) reached ∼500 mm3, they were excised from the mice. Tumors were then cut into 2×2×2 mm3 tissue explants and cultured on gelatine sponges in 12-well tissue culture plates for specific time points as indicated in the text. The DMEM culture media (Gibco) containing 20% FBS (Gibco), 1 mM sodium pyruvate (Biological Industries), 2 mM L-glutamine (Biological Industries), 1% penicillin/streptomycin/amphotericin (Biological Industries), 0.1 mM MEM nonessential amino acids (Biological Industries), 10 mM HEPES (Biological Industries), 1% BIO-MYC (Biological Industries) and 50 μg/ml gentamicin (Gibco). For drug treatment, the 2 × 2 × 2 mm^3^ explants were treated with 3 μM BAY-876 for indicated time in 37°C, 5% CO2.

### Data availability

The data supporting the findings of this study are available within the paper and its supplementary information files and are available from the corresponding authors. Code to reproduce the bioinformatics analyses and their related data is available at https://github.com/bhklab/TNBC_BAY-876.

## Acknowledgements

This study was conducted with the support of the Terry Fox Research Institute (New Frontiers Research Program PPG-1064, to D.C., B.H.K., M.L. and C.A), Canadian Cancer Research Society and the Ontario Institute for Cancer Research through funding provided by the Government of Ontario and the SGC. This work was also supported by the Canadian Institute for Health Research (CIHR project grant: Funding Reference Number 136963 to M.L., 363288 to B.H.K., CIHR FDN154328 to C.H.A. and the CEEHRC team grant: Funding Reference Number 158225 to M.L. and C.H.A.), the Princess Margaret Cancer Foundation (M.L.) and the Gattuso-Slaight Personalized Cancer Medicine Fund at Princess Margaret Cancer Centre (B.H.K.). We acknowledge the Princess Margaret Bioinformatics group for providing the infrastructure assisting us with analysis presented here. Development of PDXDTO-B64 and PDXDTO-B81 was done in collaboration with the Princess Margaret Living Biobank. We thank the patients who provided tissue, the MUHC breast surgeons and pathologists, Virginie Pilon and Anie Monast for assistance with animal studies. G.D. is a recipient of fellowships from the CIHR, of the Fonds de Recherche en Santé du Québec (FRSQ) doctoral research award and of a Cancer Research Society Next-Generation of Scientists transition award. M.L. holds an Investigator Award from the Ontario Institute for Cancer Research and the Bernard and Francine Dorval Award for Excellence from the Canadian Cancer Society. The SGC is a registered charity (number 1097737) that receives funds from AbbVie, Bayer Pharma AG, Boehringer Ingelheim, Canada Foundation for Innovation, Eshelman Institute for Innovation, Genome Canada through the Ontario Genomics Institute [OGI-055], Innovative Medicines Initiative (EU/EFPIA) [ULTRA-DD grant no. 115766], Janssen, Merck KGaA, Darmstadt, Germany, MSD, Novartis Pharma AG, Ontario Ministry of Research, Innovation and Science (MRIS), Pfizer, São Paulo Research Foundation-FAPESP, Takeda, and Wellcome.

## Author Contributions

Q.W. designed and conducted the bulk of the experiments. W.B. implemented the bulk of the computational and statistical approaches. G.D. contributed to shape the research and revised the manuscript for important intellectual content. G.D., A.-M. F., H.K.and J.C. conducted the PDXTOs assays. J.H. provided samples for PDXTEs. A.M. performed TCGA analysis. T.D.M helped histopathological image scanning. S.D contributed to interpret histopathological results of the tumor explant samples. S.D, E.L.M, N.D.A, W.J.C, S.Z. R.D. contributed to cell growing, d immunoblotting, knockdown, seahorse assays. H.M. contributed to profile proteomics data. S.C. contributed to explant work. S.J.D, M.P., D.W.C., B.H.K., M.L. and C.H.A. supervised the research. M.L. and C.H.A. oversaw the project. Figures were designed and prepared by Q.W. and W.B. The manuscript was written by Q.W, M.L. and C.H.A. with assistance from all authors. All authors reviewed the final version of the manuscript and agreed with its content and submission.

