## Supplementary material for "GLUT1 inhibition blocks growth of RB1-positive Triple Negative Breast Cancer": suplemental figures

**Supplementary Table**

| **Inhibitors** | **Target** | **Glut1 selectivity** | **Reference** |
|---|---|---|---|
| STF31 | GLUT1; NAMPT | n.d | (1,2) |
| WZB-117 | GLUTs | inhibits glucose transport and cancer cell proliferation with $IC_{50}$ of 10 µM | (3,4) |
| Fasentin | GLUT1/GLUT4; Fas-Sensitizer | preferentially inhibits GLUT4 ($IC_{50}$=68 µM) over GLUT1 | (5) |
| BAY-876 | GLUT1 | oral bioavailable GLUT1 inhibitor with $IC_{50}$ of 2 nM; displays >100-fold selectivity against GLUT2, GLUT3, and GLUT4 | (6) |

**Supplementary Table 1: List of published inhibitors targeting glucose transporters 1.** Chemical probes, their respective target names, selectivity for GLUT1 and the references.

**Supplementary Figures:**

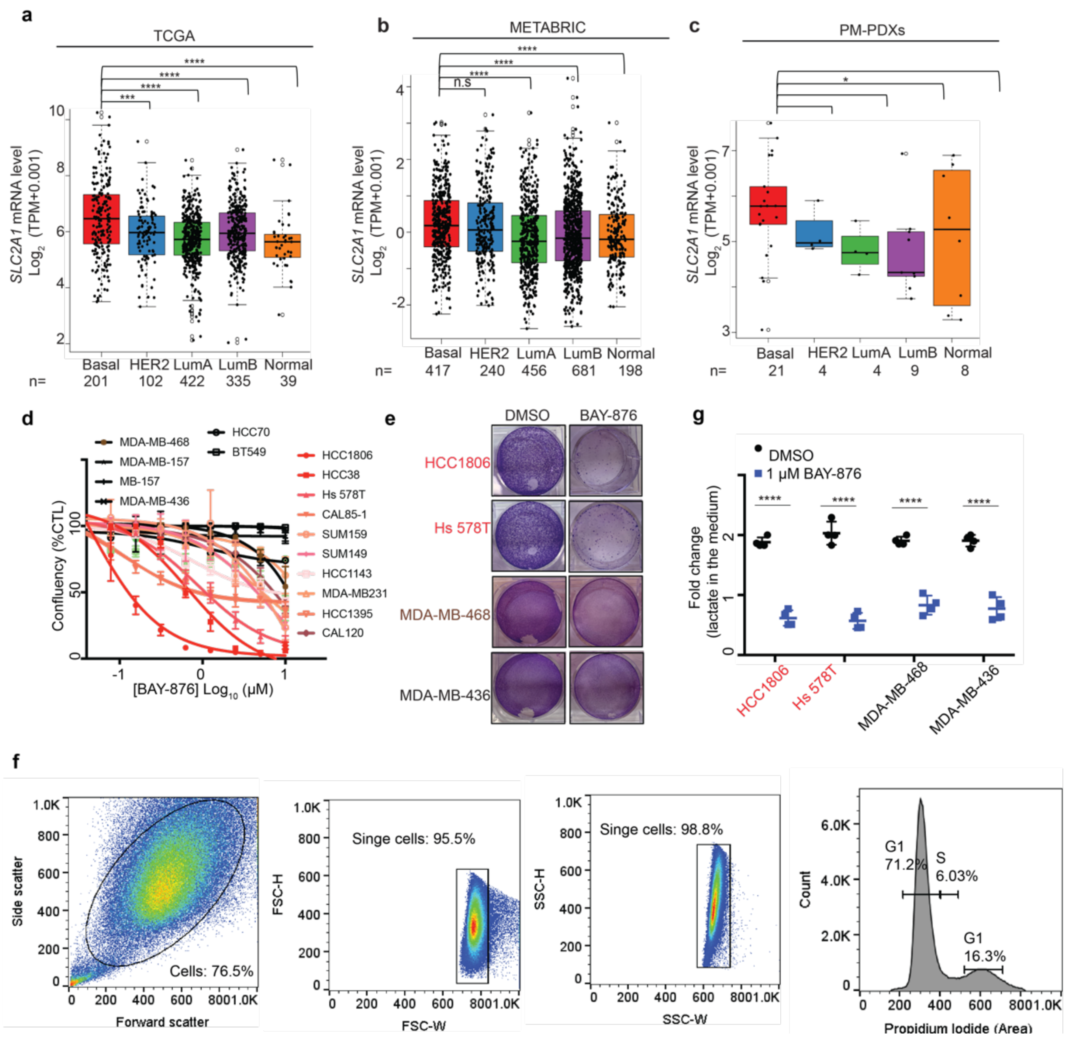


**Fig. S1**: **GLUT1 inhibition suppresses the growth of a subset of TNBC lines. (a)** *SLC2A1* gene expression in the **(a)** TCGA Breast datasets, **(b)** METABRIC Breast cancer datasets, and **(c)** Princess Margret Hospital PDXs datasets (PM-PDXs). According to PAM50 classification, the cohorts were designated as basal, HER2, LumA, LumB and normal. Gene expression is reported as $\log_2$(TPM+0.001). The number of patients (n) per group is indicated. *P* values were determined using a Wilcoxon rank sum test. *p<0.05; **p<0.01; ***p<0.001; ****p<0.0001. **(d)** Growth curves of 21 breast cancer lines with indicated concentration of BAY-876 treatment for 5 days. **(e)** Long-term colony formation assays of cell lines deemed sensitive (HCC1806 and Hs578T) and insensitive lines (MDA-MB436 and MDA-MB468) in the short-term viability assays. Cells (2000-6000 per six-well plate) were grown in the absence or presence of 1 μM BAY-876 for 14 days, stained with crystal violet and photographed. Representative plots are shown. **(f)** Representative plots for gating strategy of cell cycle analysis. **(g)** Lactate uptake analysis using Bioprofile Flex analyzer (Nova Biomedical) were performed in cells following BAY-876 treatment for the indicated time. Data shown are mean ± s.d. of n = 3 independent assays.

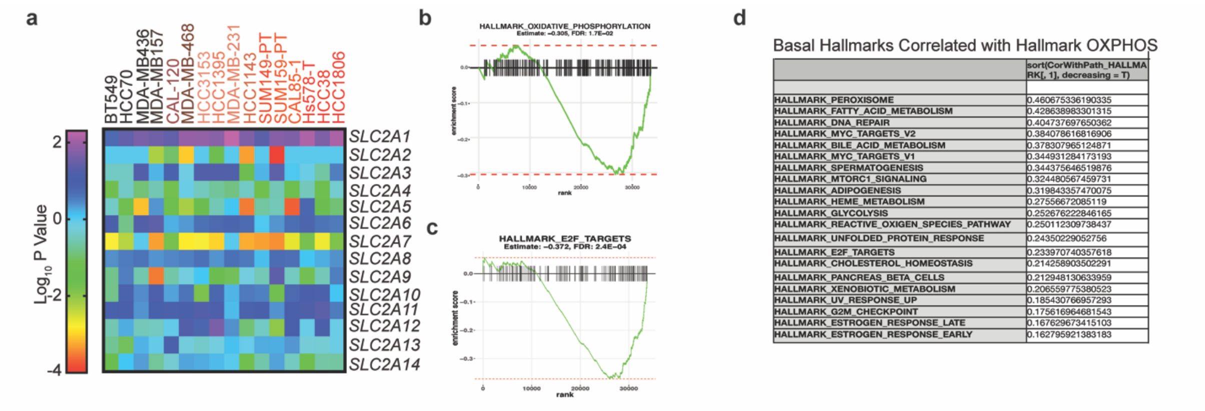


**Figure S2: RB1-E2F pathway significantly correlate with BAY-876 sensitivity. (a) Correlation** heatmap of expression level of 14 glucose transporters and BAY-876 sensitivity. Color grading of cell lines corresponds to non-responders (black) and non-responders (red) according to the $IC_{50}$ of BAY-876. **(b)** OXPHOS as a significant enriched pathway in non-responders compared to responders based on GESA on RNA-sequencing data generated. **(c)** E2F Targets as a significant enriched pathway in non-responders compared to responders based on GESA on RNA-sequencing data generated. **(d)** Top correlated pathways correlated with OXPHOS in TCGA dataset cohort.

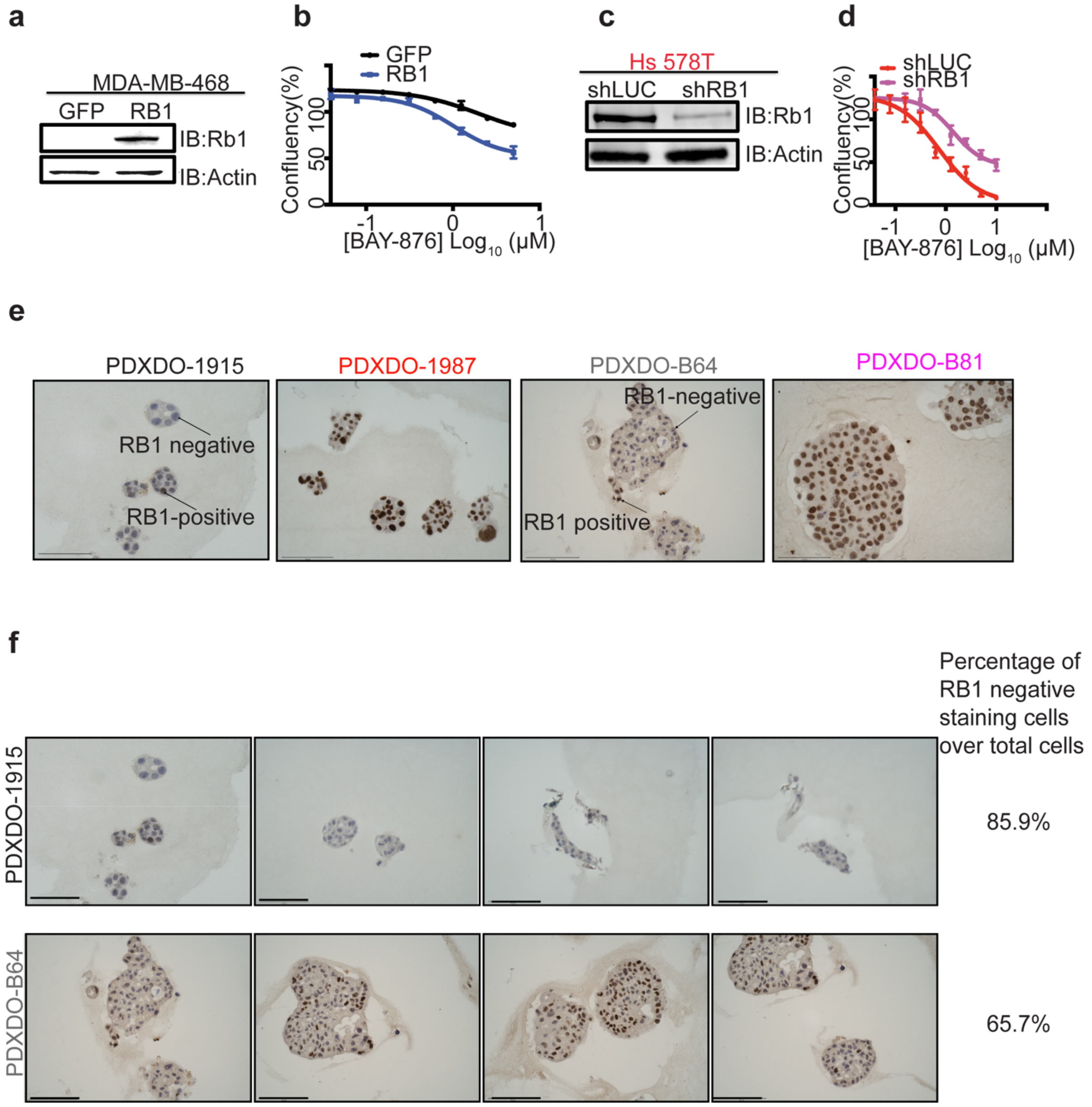


**Figure S3: RB1 levels dictates BAY876 sensitivity in TNBC lines. (a)** Representative western blot showing MDA-MB468 cells expressing RB1 or GFP control proteins. β-actin was used a loading control. **(b)** Growth curves of MDA-MB-468 cells expressing RB1 or GFP control in the presence of indicated concentrations BAY876 treatment for 5 days. Data shown are mean ± s.d of n = 4 independent experiments. **(c)** Representative western blot showing Hs578 T cells transfected with shRNA targeting RB1 or shRNA targeting control luciferase. β-actin was used a loading control. **(d)** Growth curves of Hs 578T cells with control knockdown or RB1 knockdown in the presence of indicated concentrations BAY-876 treatment for 5 days. Data shown are mean ± s.d of n = 4 independent experiments. **(e)** Representative images of RB1 IHC staining for PDXDOs. Arrows indicate different cells showing distinct staining intensity. Scale bars represent 100 μm. **(f)** Images of RB1 IHC staining for RB1 negative PDXDOs, PDXDO-1915 and PDXDO-B64. Scale bars represent 100 μm. Percentage of RB1 negative staining cells were calculated for each PDXDO.

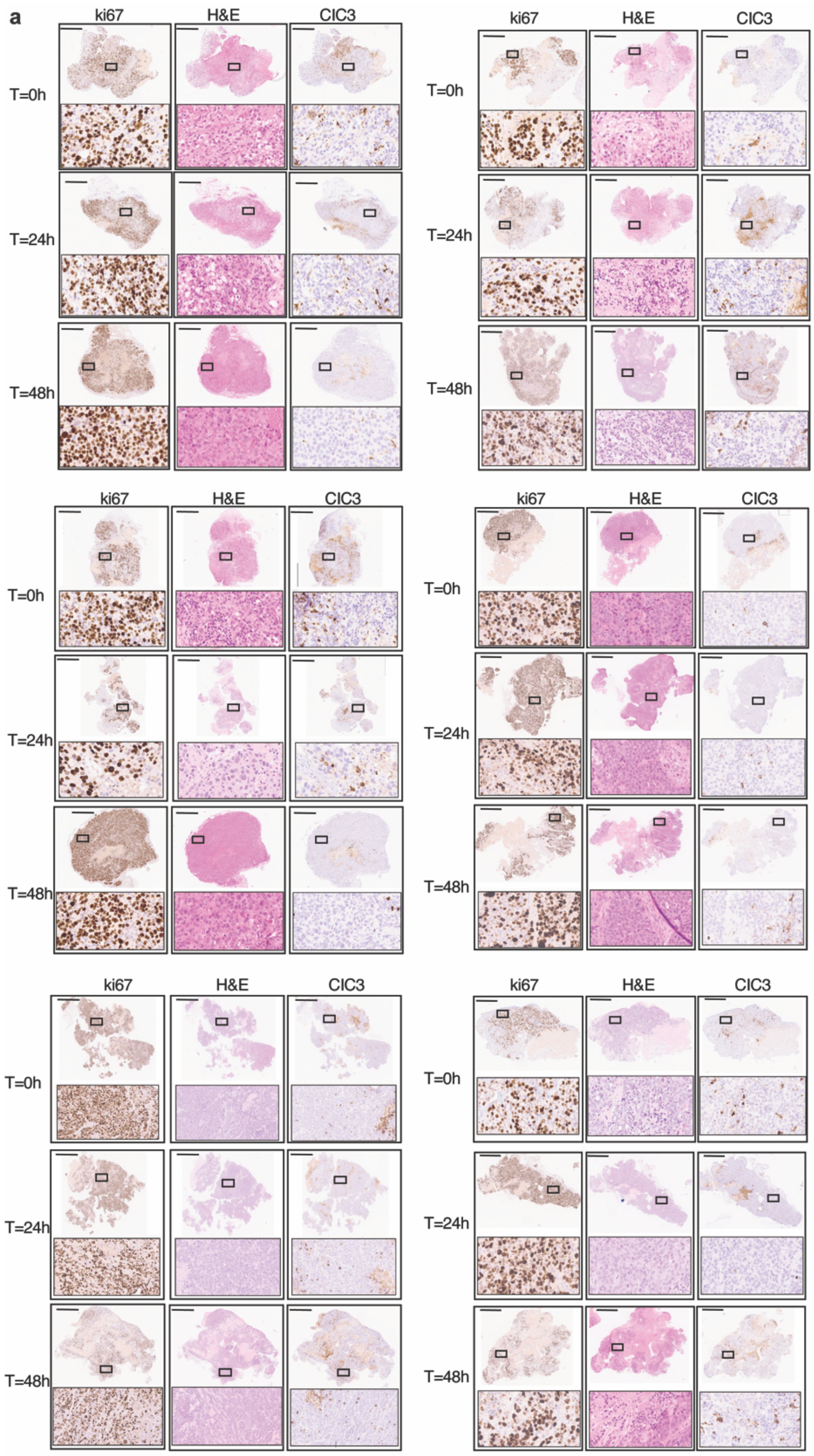


**Figure S4: Viability of explants during *ex vivo* culture time range. (a)** Viability of explants during *ex vivo* culture time range assessed by proliferative index ki67 staining, H&E staining and cleaved caspase 3 (ClC3) staining. Scale bars represent 500 µm in the representative IHC staining images. Indicated area is zoomed in 5X at the bottom.

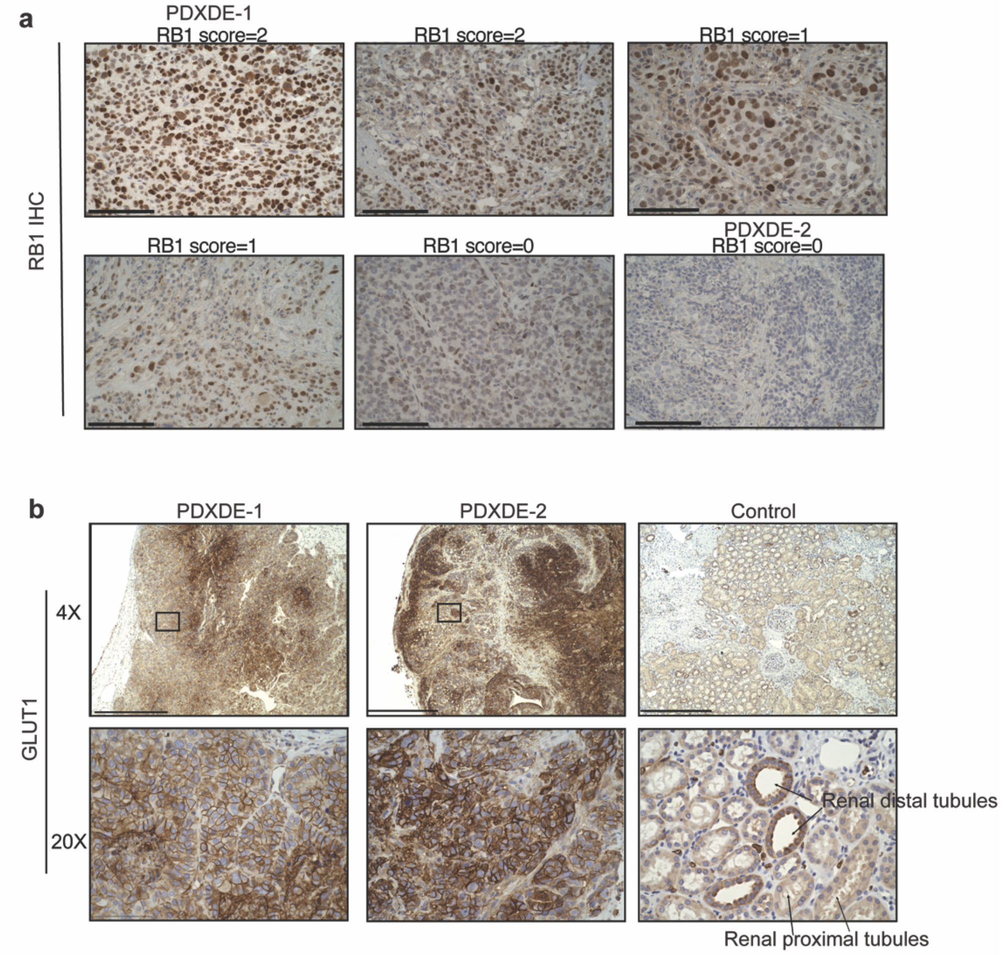


**Figure S5: On-target effect of BAY-876 in PDXDEs. (a)** Representative immunohistochemistry staining images of six PDXDEs from different TNBC patients. Scale bars represent 100 µm. RB1 IHC scores as indicated. **(b)** Representative GLUT1 IHC staining images of PDXDE-1, PDXDE-2. GLUT1 staining of human kidney tissue as negative control and positive control. Renal proximal tubules cells lacking GLUT1 expression as negative control, while renal distal tubules cells as positive control (7). Scale bars represent 500 µm. Indicated area is zoomed in 5X at the bottom.

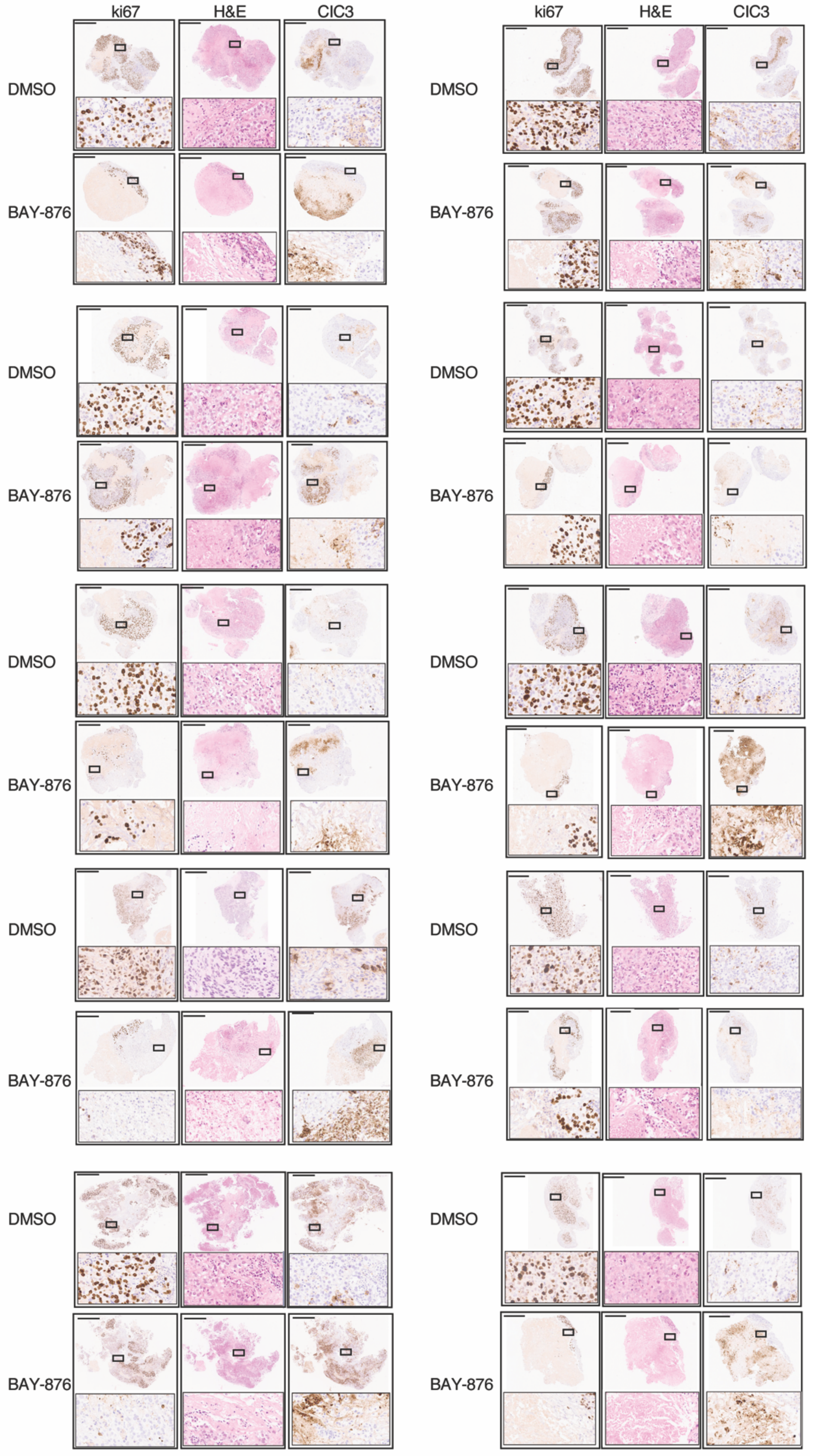


**Figure S6. IHC staining images of PDXDE-1.** Scale bar represents 500 μm. Indicated area is zoomed in 5X at the bottom.

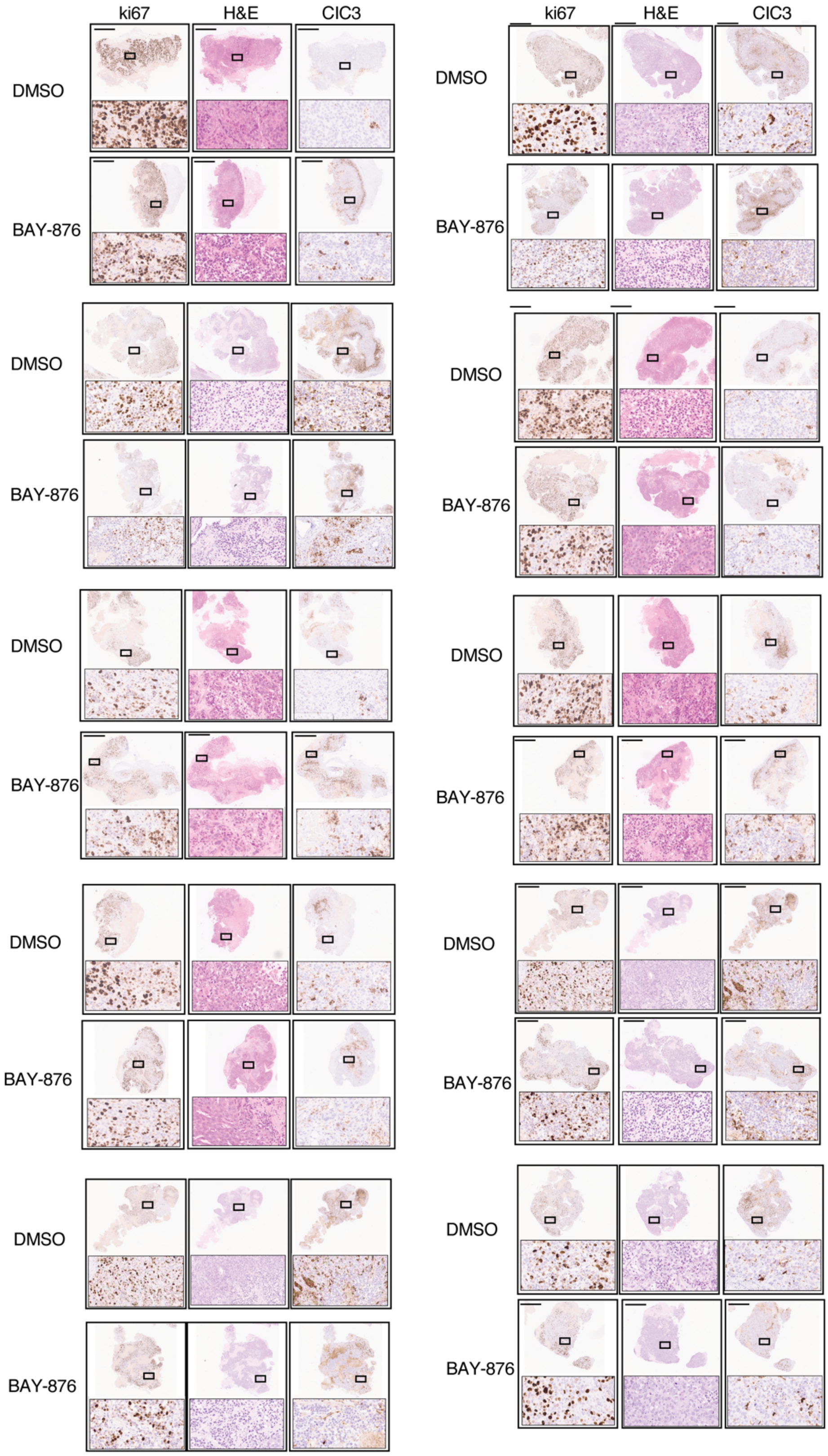


**Figure S7. IHC staining images of PDXDE-2.** Scale bar represents 500 µm. Indicated area is zoomed in 5X at the bottom.
